# Comparative analysis of biological aspects of *Leishmania infantum* isolates

**DOI:** 10.1101/2020.03.04.977462

**Authors:** Taiana Ferreira-Paes, Karen S. Charret, Merienny R.S. Ribeiro, Raquel F. Rodrigues, Leon L. Leon

## Abstract

*Leishmania infantum infantum* (LII) is one of the species that causes visceral leishmaniasis (VL) in the Old World, while *L. infantum chagasi* (LIC), and is present in the New World. Few studies address the biological differences, as well as the behaviour of these strains during infection. These parasites live inside the cells of their hosts, continuously evading the microbicidal mechanisms and modulating the immune response of these cells. One of the mechanisms used by these protozoa involves the L-arginine metabolism. Given the importance of the understanding of differences between *Leishmania* species, as well as establishing a better murine model to study leishmaniases, the objectives of this work were to analyse the biological and molecular differences between two *Leishmania infantum* strains (LII and LIC) and the degree of susceptibility of mice with different genetic backgrounds to infection, as well as to understand the role of arginase (ARG)/nitric oxide synthase (NOS) in the parasite-host relationship. The infectivity *in vivo* and *in vitro* of LII and LIC was performed in BALB/c and Swiss Webster mice, as well the NOS and ARG activities. The LII strain showed more infective than the LIC strain both *in vivo* and *in vitro*. In animals infected by both strains, a difference in NOS and ARG activities occurred. *In vitro*, promastigotes of LII isolated from BALB/c and Swiss Webster mice showed higher ARG activity than the LIC during the growth curve, however, no difference was observed in intracellular NO production by promastigotes between these strains. A comparison of the sequences of the ARG gene was made and both strains were identical. However, despite the similarity, the strains showed different expression of this gene. It can be concluded that although *L. chagasi* strains are considered identical to *L. infantum* strains, both have different biological behaviour.

## Introduction

Leishmaniases are a group of infectious diseases found worldwide and are caused by protozoa of the genus *Leishmania*. These diseases are distributed in 200 countries or territories, and approximately 20 thousand deaths each year are attributed to them, which can manifest in various forms with different symptoms depending on the infecting species and the host immune response. The main forms of the disease are cutaneous, mucocutaneous and visceral leishmaniases (VL), the latter being the most lethal manifestation with almost 0.5 million new cases each year, if not treated the mortality rate can reach 100% in two years ^[1,2]^.

*Leishmania infantum* is one of the species that causes VL and is present in the Mediterranean Basin, as well as in the Middle East and South Asia. Some years ago, *L. chagasi* was considered a new species of *Leishmania* causing visceral leishmaniasis in the New World. However, after genetic sequencing studies, *L. chagasi* was considered identical to *L. infantum*, being denominated *L. infantum* (syn. *L. chagasi*) ^[3]^. Nevertheless, further studies showing antigenic differences between both strains, they have considered as two subspecies *L. infantum infantum* in the Old World and *L. infantum chagasi* in the New World ^[4]^.

Our previous studies have reported a possible relationship between infectivity and the metabolic pathway of L-arginine not only in *L. infantum chagasi* but also in *L. amazonensis*, and *L. braziliensis* ^[5-7]^. L-Arginine is metabolized mainly by nitric oxide synthase (NOS) and arginase (ARG) ^[8]^, and the amount of this amino acid accessible is critical for *Leishmania* proliferation ^[9,10]^. NOS and ARG can act directly in the intracellular fate of the parasite in macrophages; NOS activity results in the production of nitric oxide (NO), which is harmful for the parasite, whereas ARG produces L-ornithine, essential to parasite proliferation ^[11-14]^.

In experimental chemotherapy studies, the choice of the animal model is a primordial issue. Nowadays, for VL, mice are usually the chosen models, due to the easiness of manipulation and their similarity with the human genome ^[15]^ and different lineages are used aiming to mimic human leishmaniasis manifestations ^[16]^. Resistance/susceptibility to *Leishmania major* infection has been associated with the balance of Th1 and Th2 immune responses ^[17-19]^; however, for other *Leishmania* species, such profile is not so clear-cut, hampering the study of the pathophysiology of the disease and the development of novel drugs. BALB/c mice (inbred) are highly susceptible to infection by *Leishmania* spp., since they present Th2 immune response and develop chronic disease a few weeks post-infection. In addition, due to the genetic variation of the human population, the infections are quite heterogenous, which makes it difficult to trace similarities with infections in inbred mice. In accordance with the above and based on our observations, the aim of this work was to demonstrate biological differences between these two *L. infantum* strains *in vivo* and *in vitro*, the susceptibilities of mice with different genetic backgrounds to infection by these parasites, as well as to understand the role of arginase (ARG)/nitric oxide synthase (NOS) in the parasite-host relationship.

## Materials and methods

### Parasites

Two strains of parasites were used: *Leishmania infantum infantum* (MHOM/MA/67/ITMAP263; from now called LII) and *Leishmania infantum chagasi* (MCAN/BR/97/P142; from now called LIC). In all protocols, infective isolates newly isolated from BALB/c and Swiss Webster infected mice were used. The promastigote forms were cultured at 26°C in Schneider’s Insect Medium (Sigma Aldrich, St. Louis, MO, USA) supplemented with 20% foetal calf serum (FCS), 100 U/mL Penicillin G potassium and 100 µg/mL streptomycin.

### Mice infection

Female BALB/c and Swiss Webster mice (6-8 weeks of age) were obtained from the Instituto de Ciência e Tecnologia em Biomodelos (ICTB, FIOCRUZ, Rio de Janeiro, Brazil). All procedures involving animals were approved by the Ethics Committee on the Use of Animals at FIOCRUZ (CEUA LW-026/15). Six groups of animals (5 mice/group) were infected or not by an intraperitoneal injection of 1 × 10^8^/100µL parasites as follows: Group 1: non-infected BALB/c mice (Control); Group 2: BALB/c mice infected by LIC; Group 3: BALB/c mice infected by LII; Group 4: non-infected Swiss Webster mice (Control); Group 5: Swiss Webster mice infected by LIC; and Group 6: Swiss Webster mice infected by LII. The animals were maintained for 30- and 60-days post-infection (dpi), when they were euthanized.

### Limiting dilution analysis (LDA)

Parasite quantification in the spleen and liver was performed by the limiting dilution method. Mice were euthanized at 30 and 60 dpi, the organs were aseptically removed, weighted and homogenized in Schneider’s medium. The cells were centrifuged at 1,500 rpm for 10 min, and 2 × 10^5^ cells/well were placed in a 96-well plate containing Schneider’s medium plus 20% FCS using a serial dilution (1:10). For 7 days, the plates were maintained at 26°C and the wells were daily examined using an inverted microscope. The number of parasites/mg of tissue was estimated based on the total weight of the removed tissue and the parasite load in the serial dilution according to Taswell ^[20,21]^.

### NOS activity in spleen and liver cultures

The NO present in the supernatant of cultures was evaluated indirectly by Green’s method ^[22]^. Briefly, cells from the infected organs were maintained in culture for 48 h. Afterward, 100 µL supernatant were collected and mixed (v/v) with Griess reagent (0.1% N-1-naphtyl-ethylenediamine dihydrochloride in a solution of 5% phosphoric acid and 1% sulphanilamide). After 10 min at room temperature, the NO production was measured at 540 nm, using sodium nitrite in Schneider’s Insect Medium, in a concentration range of 0.78 to 200 µM, as standard.

### ARG activity in spleen and liver cultures

Infected organ cells (1 × 10^7^) were treated as previously described by Corraliza and collaborators (1994) ^[23]^. Briefly, spleen and liver cells were previously washed in sucrose and KCL solution and an anti-proteolytic buffer was added. After cell lysis, L-arginine (0.5M) at pH 9.7 was added. The samples were incubated at 37°C and the reaction was stopped by the addition of an acidic solution (H_2_SO_4_, H_3_PO_4_ and water). The amount of urea produced was measured by adding 25 µL 9% α-isonitrosopropiophenone in ethanol and heating at 100°C for 45 min. After 10 min in the dark, the absorbance was determined at 540 nm, using a urea solution in Schneider’s Insect Medium, in a concentration range of 1.5 to 30 µg/mL, as standard.

### Promastigote proliferation

Promastigotes were obtained from amastigotes freshly isolated from the four infected experimental groups named LIC.B (*L. infantum chagasi* isolated from BALB/c mice); LII.B (*L. infantum infantum* isolated from BALB/c mice); LIC.S *(L. infantum chagasi* isolated from Swiss Webster); LII.S *(L. infantum infantum* isolated from Swiss Webster). The promastigotes were cultured as previously described. The initial inoculum used was 5 × 10^5^ promastigotes/mL and the cultures were counted daily for up to 7 days, cells and supernatant were collected for use in the determination of the percent of metacyclogenesis, the infectivity to macrophages, the activity of NOS and ARG gene sequencing and expression.

### Metacyclogenesis of promastigotes

To determine the percentage of metacyclic parasites of groups LIC.B, LII.B, LIC.S and LII.S during *in vitro* proliferation, a complement lysis test was performed. At 48, 72 and 96 h, the parasites were collected, washed in PBS, the suspension adjusted to 1 × 10^6^ parasites/mL in PBS and was incubated with 20% human complement (Sigma Aldrich). After 30 min, the number of resistant parasites was counted and the percentage of metacyclic cells was calculated.

### Infectivity to peritoneal macrophages

Peritoneal macrophages were removed from BALB/c and Swiss Webster mice to evaluate the infectivity of the four isolates of both *Leishmania* strains (LIC.B, LII.S, LIC.B and LIC.S). The peritoneal cavity of mice was washed with RPMI 1640 medium (Sigma Aldrich) supplemented with 10% FCS. The cells were adjusted to 2 × 10^5^ macrophages/well, placed in a 96-well plate and maintained for 1 h at 37°C with 5% CO_2_. Parasites were added to the cell cultures at a ratio of 5:1 and maintained overnight, followed by the removal of non-internalized parasites. Cell cultures were kept for 24 and 72 h and stained with Fast Panotic (Laborclin, Pinhais, PR, Brasil). The infection index was calculated by the following formula:

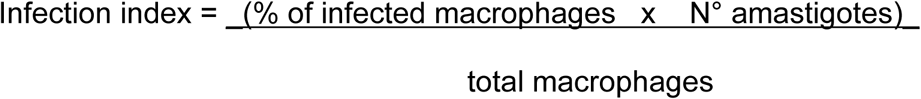

### NOS and ARG activities in promastigotes

To evaluate the production of NO within promastigotes (LIC.B, LII.S, LIC.B and LIC.S), the parasites (2 × 10^6^/mL) were incubated with 200 µL L-arginine solution and 5 µM DAF-2DA (Sigma) for 2 h at room temperature. The resulting fluorescent compound was measured by fluorimetry with an emission wavelength at 485 nm and excitation wavelength at 530 nm. The assay specificity was confirmed using RAW 264.7 cells as a positive control. To analyse the ARG activity, promastigotes (1 × 10^7^) cultivated in Schneider’s medium plus 20% FCS were harvested at 48, 72 and 96h. The amount of urea produced by them was measured as previously described.

### RNA purification, cDNA synthesis and qPCR of ARG

Total RNA was extracted from 10^7^ promastigotes of the four isolates, LIC.B, LII.S, LIC.B and LIC.S, using Trizol reagent (Invitrogen, Carlsbad, CA, EUA), following the manufacturer’s protocol. RNA (2 µg) was reverse transcribed through the GoScript™ Reverse Transcription System performed with Oligo(dT)_15_ (Promega, Madison, Wisconsin, EUA), according to the manufacturer’s instructions. The resultant cDNA was quantified in a Qubit 2.0 Fluorimeter (Thermo Fischer) and kept at - 20°C. Gene expression by qPCR was performed using a ViiA™ 7 Real-Time PCR System (Applied Biosystems, Carlsbad, CA, USA). All reactions were performed as biological triplicates and technical duplicates using a GoTaq® qPCR Master Mix Kit (Promega). The primers were designed according to the ARG gene sequence (LinJ.35.1490) obtained from the TriTrypDB.org platform (ARG: PF – 5’ GTGTGGTACGGTCTCCGGTA 3’; PR – 5’ GTGTGGTACGGTCTCCGGTA 3’). Alpha-tubulin and GAPDH were used as housekeeper genes (Alpha-tubulin: PF – 5’ CAGGTGGTGTCGTCTCTGAC 3’; PR – 5’ TAGCTCGTCAGCACGAAGTG 3’). Fluorescence readings were performed under the following conditions: pre-incubation at 95oC for 10 min, 40 cycles of amplification at 95°C for 30 s, and 60°C for 60 s. Gene expression analyses were performed using QuantStudio™ Software V1.2 (Applied Biosystems) based on ΔCt methods.

### Sequencing of the ARG gene

The ARG gene was amplified by conventional PCR from promastigote DNA. The DNA was extracted LIC.B, LII.S, LIC.B and LIC.S using the Genomic DNA Purification Kit protocol (Thermo Fisher). The conventional PCR was carried out with 50 ng of DNA, GoTaq® Hot Start Master Mixes 2x (Promega), specific primers (5’ CGCATATGATGGAGCACGTGCA 3’ and 5’ CGGGATCCCTACAGTTTGGCG) and DEPC-treated water up to 25 mL. PCR amplification was performed with a programmable thermal cycler (Applied Biosystems). The amplification protocol was carried out as follows: 1 cycle 95°C; 30 cycles of 30 s at 94°C, 30 s at 58°C, and 40 s at 72°C; and 1 cycle of 5 min at 72°C. Next, 200 ng of the PCR product and forward primer for the studied gene (described above) was used for sequencing using a Sequence Scanner (Applied Biosystems). The results were analysed using the BioEdit program (Ibis Biosciences, Carlsbad, CA, USA).

### Statistical analysis

The means and standard deviations were determined from at least three independent experiments. Statistical analyses were performed with the program GraphPad Prism 5 (GraphPad Software, USA). The ANOVA test was applied, and p values less than 0.05 were considered significant.

## Results

### Parasite load

Initially, BALB/c and Swiss Webster mice were infected (i.p.) with 1 × 10^8^/100 µL promastigotes of *L. infantum infantum* (LII) and *L. infantum chagasi* (LIC). Mice were kept for 30- and 60-days post-infection (dpi) until euthanasia. Quantifying the total spleen weight, it was observed for BALB/c mice at 30 and 60 dpi no alteration by infection with LIC or LII, while for Swiss Webster, at 60 dpi, for both strains, the spleen weight was significantly (p ≤ 0.05) higher than in non-infected counterparts (Fig 1A). No statistical difference was observed in the liver weight between non-infected and corresponding infected groups. The LDA of the spleen and liver cells of infected mice showed a significant difference in the parasitic load among the different infected groups. In the spleen, this parameter was significantly higher at 60 dpi than at 30 dpi in three experimental groups, except in Swiss Webster mice infected by LIC (Fig 1B). Comparing LII and LIC, the former was always more infective in both mice lineages, such difference being of about 3-fold in BALB/c mice at 60 dpi. In the liver, although with lower amounts of parasite than in the spleen, also higher parasite was detected in the case of LII infection (Fig. S1A).

**Figure 1.**
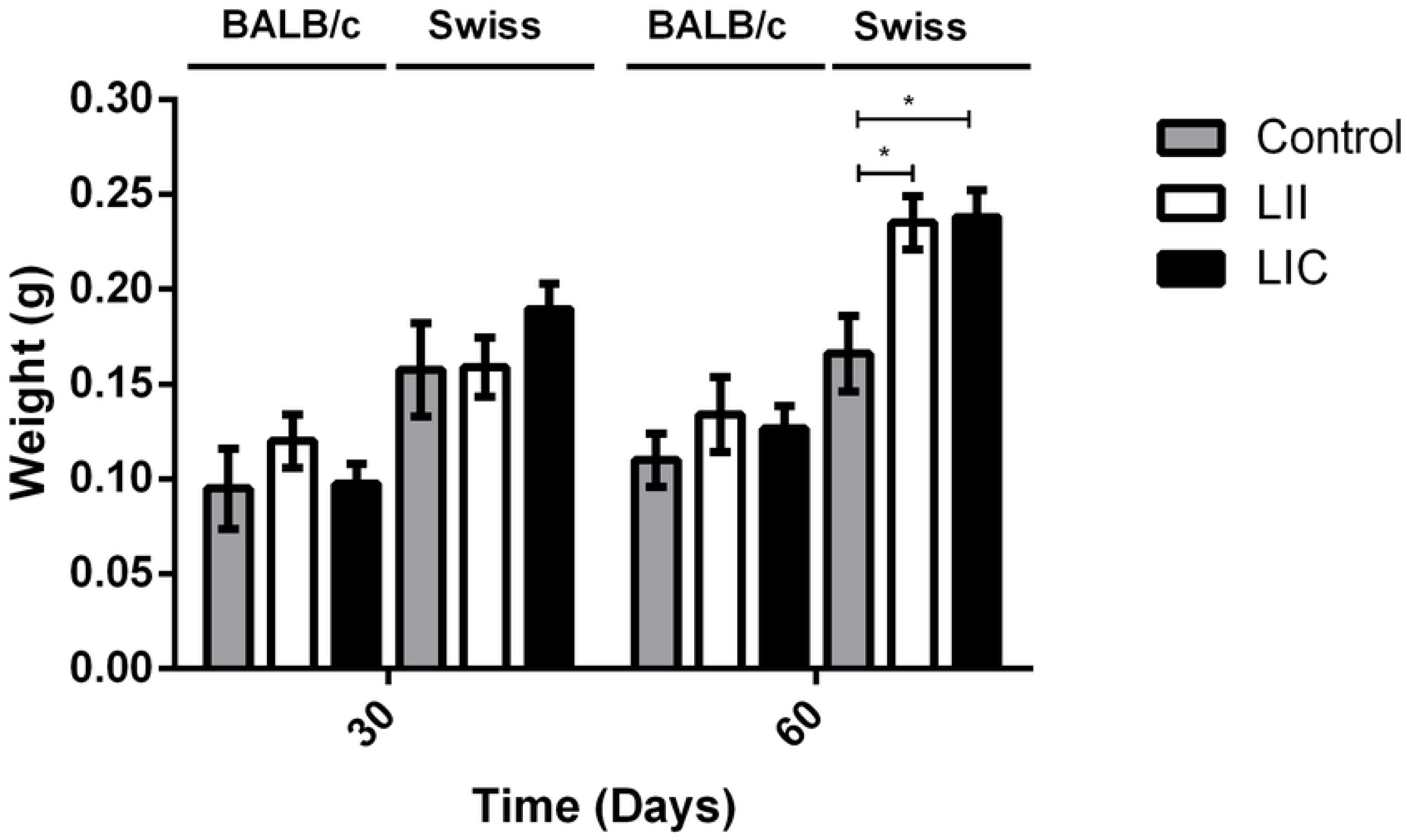
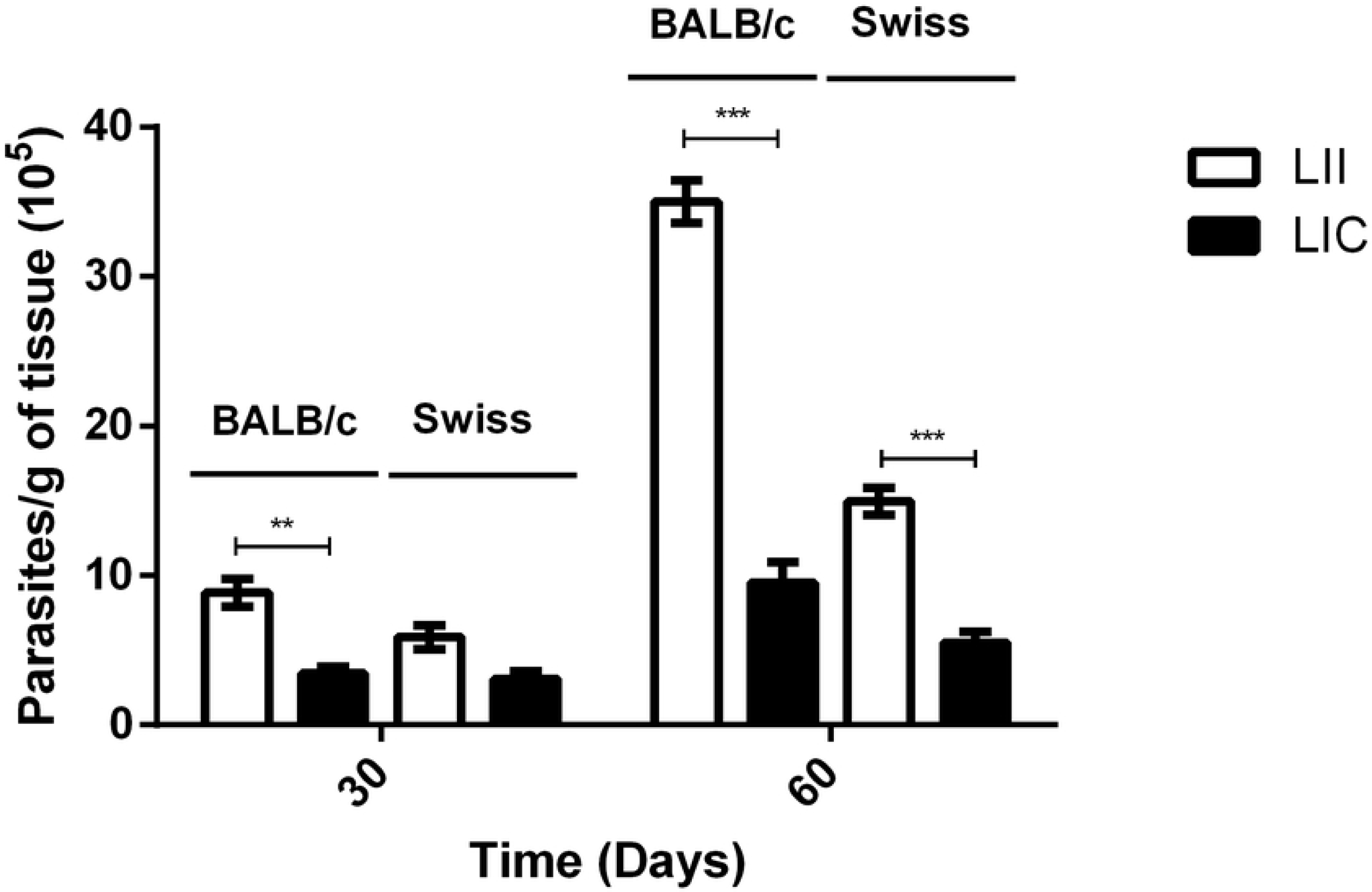
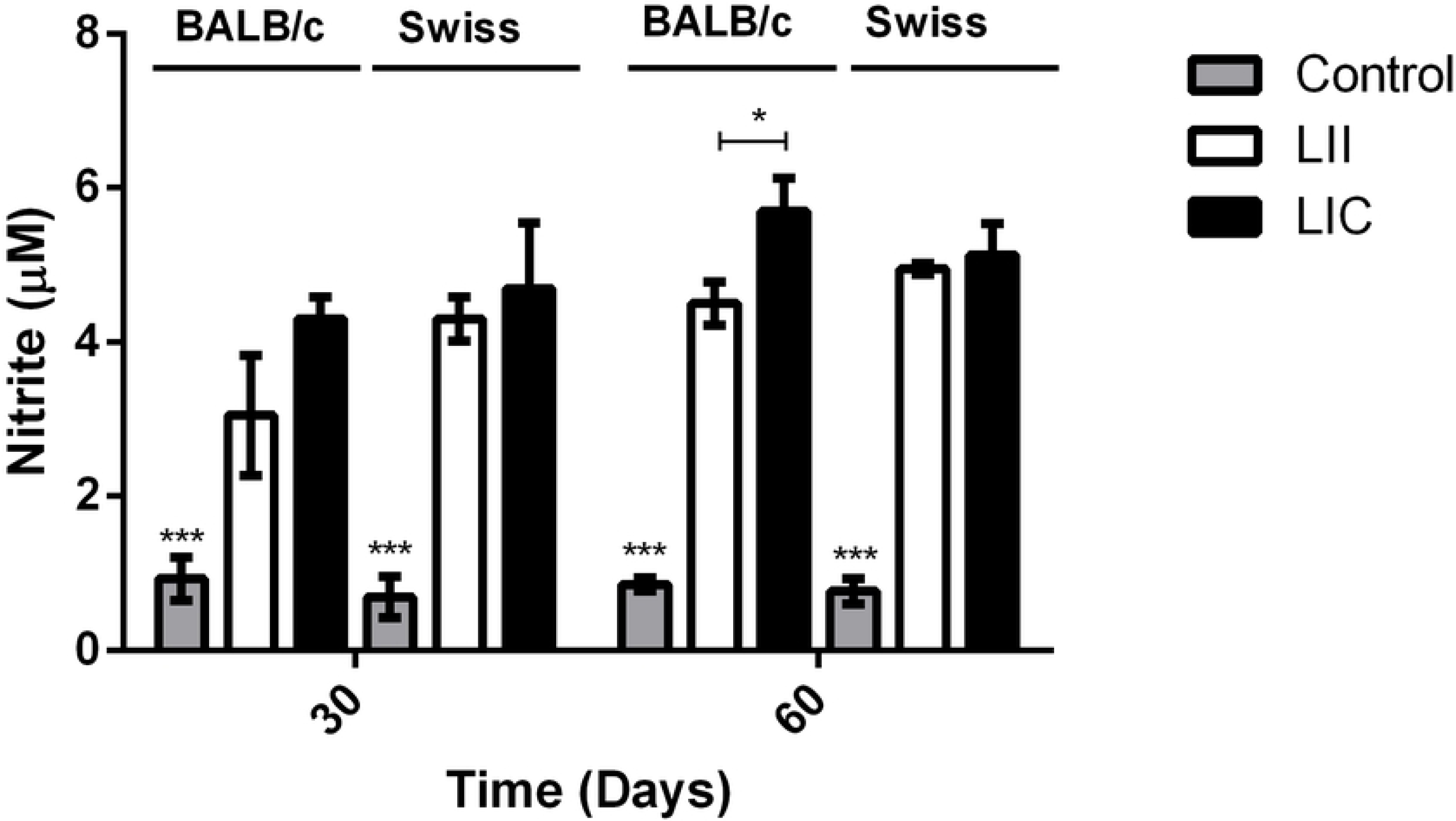
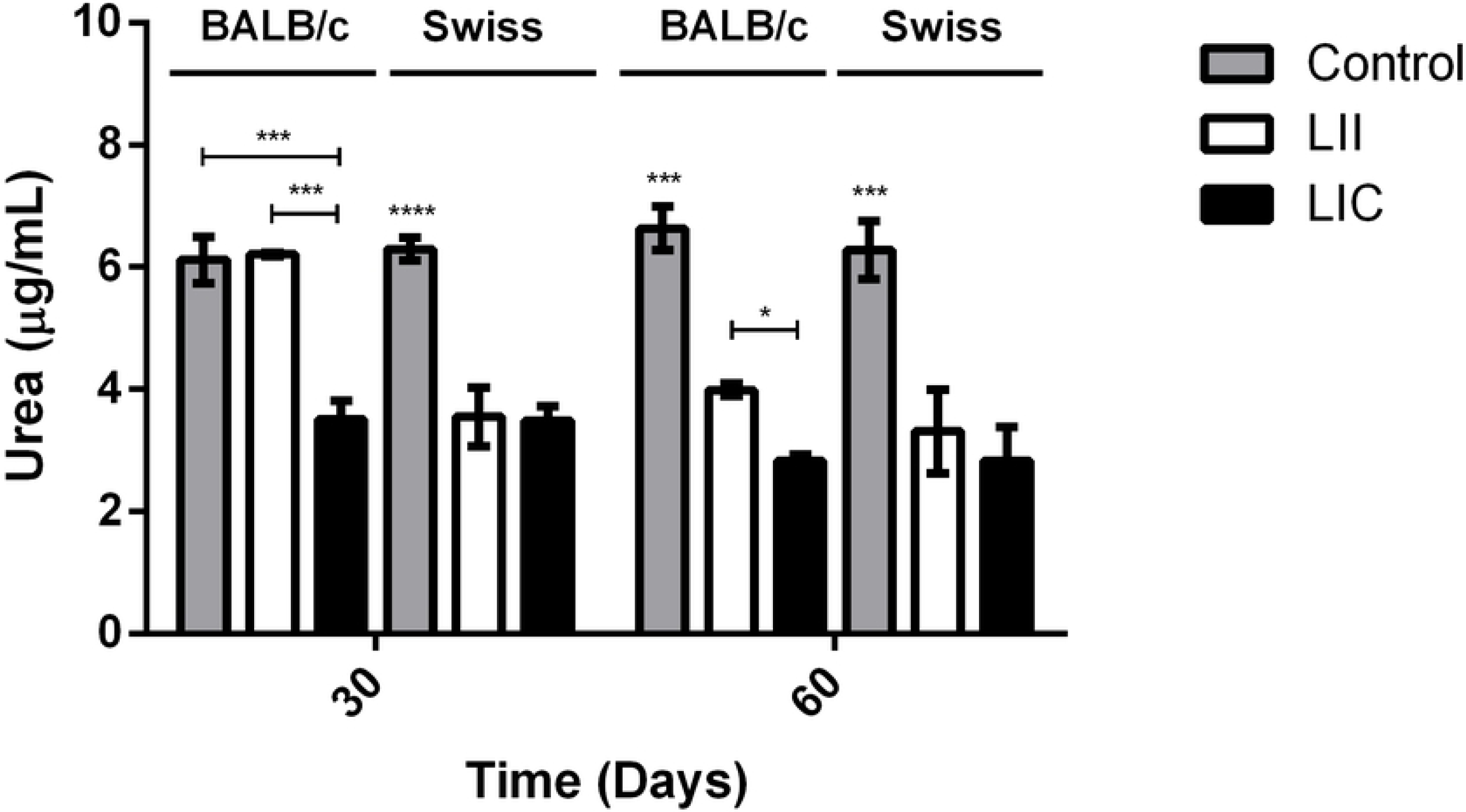
Evaluation of *in vivo* infection of two *L. infantum* strains. (A) Spleen weights of BALB/c and Swiss Webster mice infected by *L. infantum strains.* (B) Parasite load in the spleen of infected mice. (C) Nitrite levels (NOS activity) of spleen cell cultures. (D) Urea levels (ARG activity) of spleen cell cultures. The BALB/c and Swiss Webster mice infection was maintained at 30 and 30 dpi. The number of parasites/g of spleen was estimated based on total weight of the spleen removed and the parasite load in the serial dilution. The nitrite and urea levels were measured by spectrophotometry at 540 nm. Control: non-infected mice; LII (*L. infantum infantum*); LIC (*L. infantum chagasi*). *p ≤ 0.05; **p ≤ 0.0009; ***p < 0.0001

### *In vivo* NOS and ARG activities

Since the enzymes NOS and ARG are regulated by cytokines related to Th1 and Th2 type immunological response profiles, their activities may indicate the susceptibility/resistance to infection in different murine models ^[26]^. After 48 h of incubation of the spleen and liver cell cultures of the 6 groups at both 30 and 60 dpi, the supernatants were used to indirectly quantify the activities of NOS, by nitrite levels, and ARG, by urea production. As expected, there was a significant increase (p < 0.0001) in nitrite levels due to *Leishmania* infection at both 30 and 60 dpi. For BALB/c mice, higher NO levels in spleen cultures were observed after infection with LIC when compared to LII, being the differences for the two parasite strains statistically significant (p ≤ 0.05) at 60 dpi (Fig 1C), while for Swiss Webster mice, no significant differences between the infections with LIC and LII. In liver cultures from BALB/c, infection by LIC showed higher nitrite levels than those infected by LII at both 30 and 60 dpi. For Swiss Webster mice, was observed significant differences (p ≤ 0.009) only at 60 dpi, when LIC infection showed higher nitrite levels compared to LII one (Fig S1B).

Analysis of ARG in spleens from both mice lineages, revealed an overall decrease in the enzyme activity compared to non-infected controls. For BALB/c mice infected by LIC at 30 dpi, there was a significant decrease (p < 0.0001) in ARG activity in comparison with the corresponding non-infected control (Fig 1D), while such decrease was not observed in those animals infected by LII. Actually, mice infected by LII showed higher ARG activity (p < 0.0001) than those infected by LIC. However, at 60 dpi, a decrease in ARG activity (p ≤ 0.0001) was observed in BALB/c mice infected with LIC or LII when compared with non-infected mice. Nevertheless, mice infected by LII still showed more ARG activity (p ≤ 0.05) than those infected by LIC. In Swiss Webster mice, the ARG activity was significantly smaller in groups infected by LIC and LII than in the uninfected control at both 30 and 60 dpi. Also, no differences were observed between the two infected groups. Interestingly, assays with liver cell cultures, at both 30 and 60 dpi, significant decrease (p ≤ 0.0001) in ARG was observed in both mice lineages infected by LIC or LII (Fig S1C).

### *In vitro* biological behaviour of LII and LIC

Since the *L. infantum* strains (*L. infantum infantum* and *L. infantum chagasi*) showed differences in the infectivity in the two murine lineages evaluated, our next step was to analyse their *in vitro* behaviour and the potential influence of the host, through promastigotes proliferation and metacyclogenesis. Firstly, promastigotes were obtained from amastigotes from the four infected groups generating the isolates: LIC.B (*L. infantum chagasi* isolated from BALB/c mice); LII.B (*L. infantum infantum* isolated from BALB/c mice); LIC.S *(L. infantum chagasi* isolated from Swiss Webster); LII.S *(L. infantum infantum* isolated from Swiss Webster). In general, the proliferation profiles of the two *Leishmania* strains were similar, since the late log phase was between the 3^rd^ and 4^th^ days of growth. However, the parasite concentration of LII (LII.B and LII.S) was higher than that of LIC (LIC.B and LIC.S), with significant differences from the 2^nd^ to 6^th^ days of culture (Fig 2).

**Figure 2.**
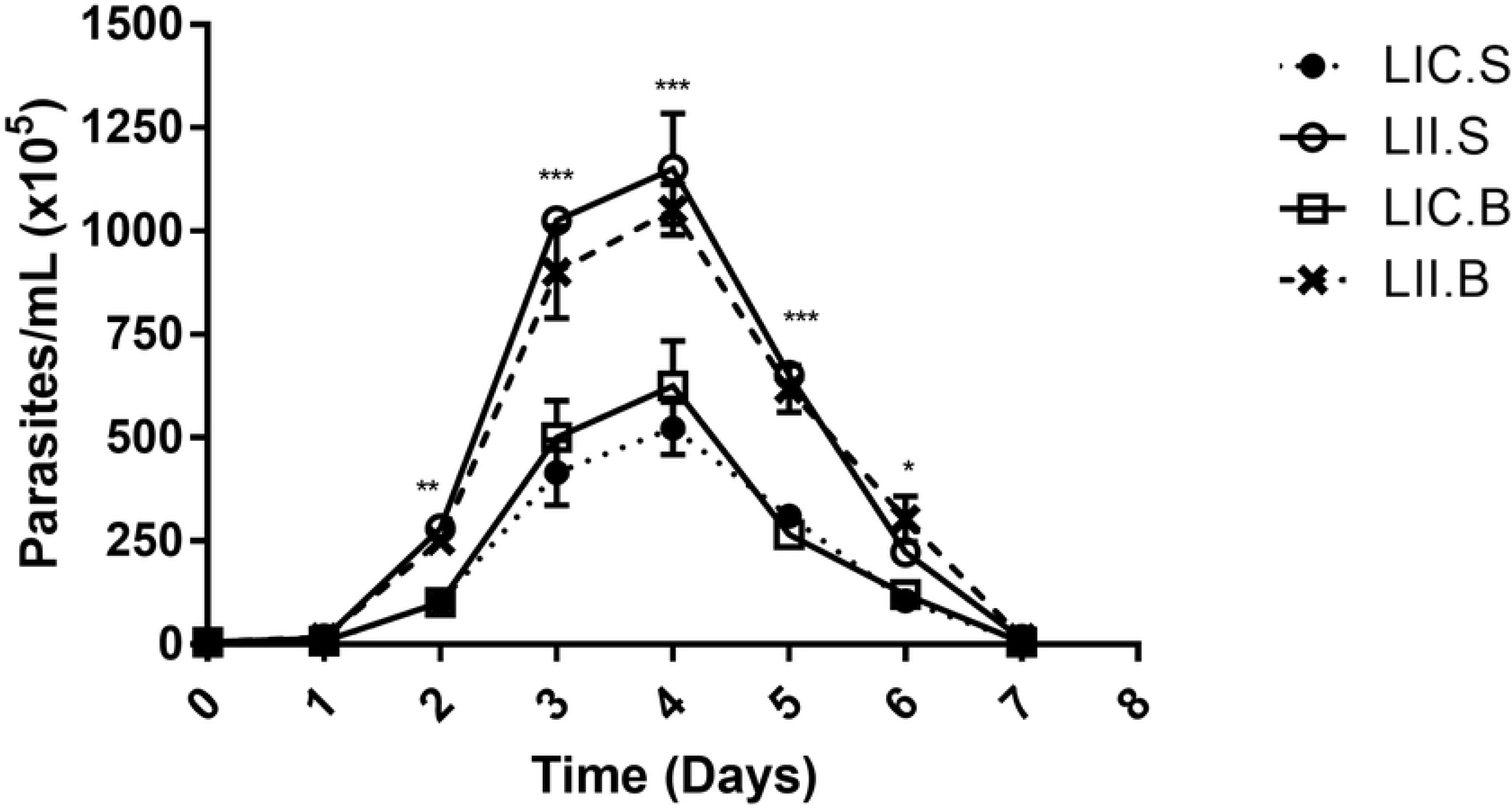
*In vitro* proliferation of promastigote forms. Promastigotes were obtained from amastigotes freshly isolated from the four infected experimental groups and cultivated for 7 days. The parasites were daily counted, and the growth profile was evaluated. LIC.B (*L. infantum chagasi* isolated from BALB/c mice); LII.B (*L. infantum infantum* isolated from BALB/c mice); LIC.S *(L. infantum chagasi* isolated from Swiss Webster); LII.S *(L. infantum infantum* isolated from Swiss Webster). *p ≤ 0.05; **p ≤ 0.009; ***p < 0.0001

Although the four isolates presented the same growth profile, their proliferation rates were different. So, the next step was to evaluate whether these parasites present differences in infectivity, using the complement lysis test in different times (48, 72 and 96 h) to define the percentage of metacyclic forms in culture. At all times evaluated, the percentage of resistant cells to complement was significantly higher in *L. infantum infantum* isolated from both BALB/c and Swiss Webster mice, suggesting its higher infectivity when compared to *L. infantum chagasi*, independently of the mice lineage from which it was isolated (Table 1). It is important to notice that the higher percentage of metacyclic forms in all different situations were at 72h, where the highest parasite concentration was observed in the growth curve.

**Table 1.** Evaluation of the metacyclogenesis for the four *Leishmania* isolates.

| % metacyclic parasites |  |  |  |  |
| --- | --- | --- | --- | --- |
| Time | BALB/c |  | Swiss Webster |  |
|  | LII.B | LIC.B | LII.S | LIC.S |
| **48h** | 56.0 ± 5.7** | 27.0 ± 3.0 | 49.0 ± 7.8* | 22.0 ± 3.5 |
| **72h** | 75.0 ± 3.5** | 38.0 ± 2.0 | 68.0 ± 4.2** | 34.0 ± 6.4 |
| **96h** | 56.0 ± 1.4* | 31.0 ± 1.4 | 43.0 ± 6.4* | 18.0 ± 1.4 |

Percentual of metacyclogenesis was determined by complement lysis test. LIC.B (*L. infantum chagasi* isolated from BALB/c mice); LII.B (*L. infantum infantum* isolated from BALB/c mice); LIC.S (*L. infantum chagasi* isolated from Swiss Webster); LII.S (*L. infantum infantum* isolated from Swiss Webster).
\*p ≤ 0.009; \*\*p < 0.0001. Comparison of metacyclic forms percentage observed on LII and LIC isolates from each mice lineage.

To confirm the hypothesis that LII is more infective than LIC, the four parasite isolates from BALB/c and Swiss mice, were used to infect peritoneal macrophages obtained from the same mice they were isolated from (Fig 3) and after 24 and 72 h, the infection index was calculated.

**Figure 3.**
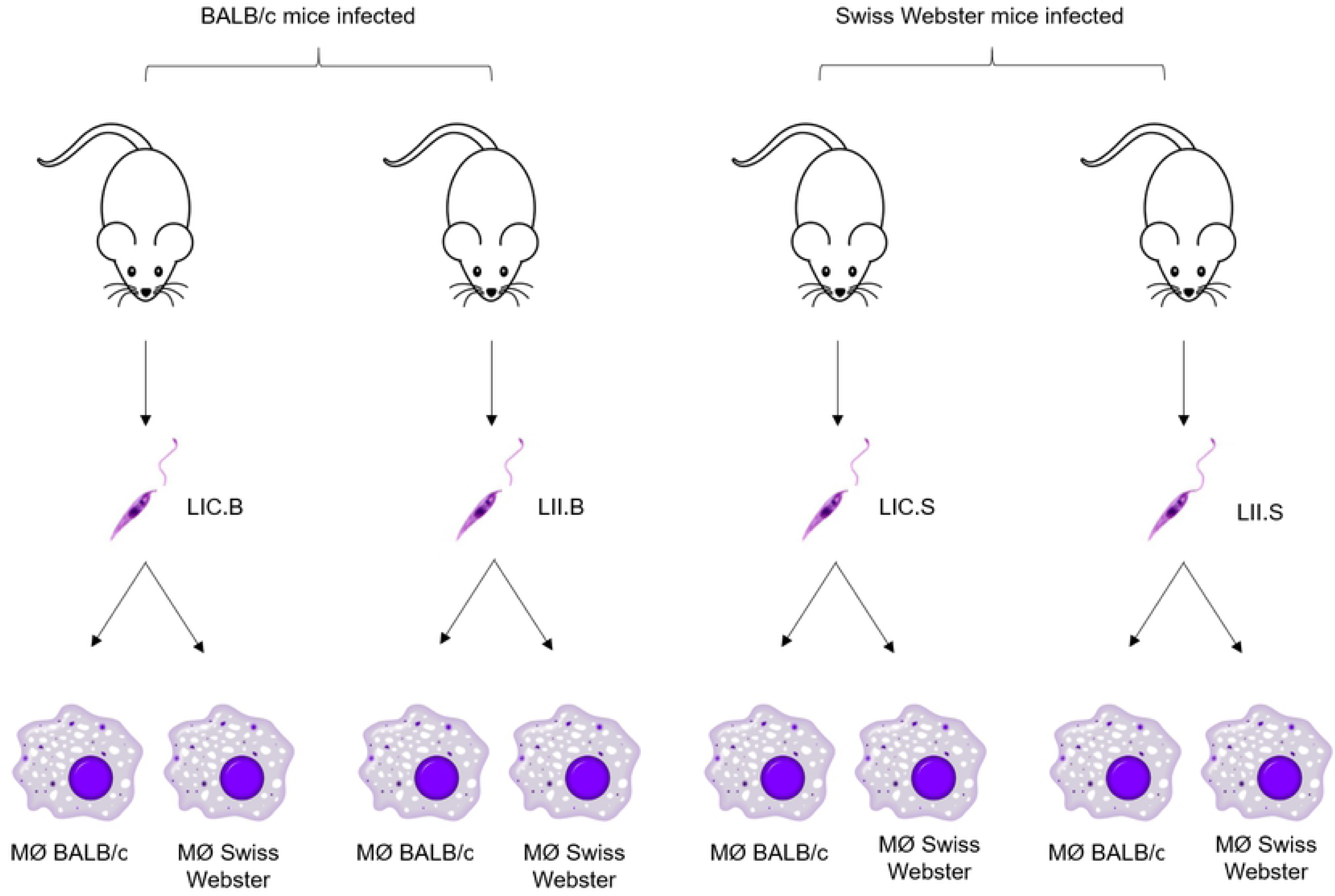
Scheme of *in vitro* infection. Four *L. infantum* isolates were used to infected peritoneal macrophages from BALB/c and Swiss Webster mice, at a ratio of 5:1. The infections were maintained for 24 and 72h. MØ (macrophage); LIC.B (*L. infantum chagasi* isolated from BALB/c mice); LII.B (*L. infantum infantum* isolated from BALB/c mice); LIC.S *(L. infantum chagasi* isolated from Swiss Webster); LII.S *(L. infantum infantum* isolated from Swiss Webster).

As expected, the values of a given combination were higher at 72 h than at 24 h. In both mice lineages (BALB/c and Swiss Webster), LII was more infective than LIC (p < 0.0001), but this infection was prominent in BALB/c mice, which showed the highest infection index at both 24 and 72 h (Table 2). This data reinforced the one obtained in both, metacyclogenesis and *in vivo* experiments.

**Table 2.**
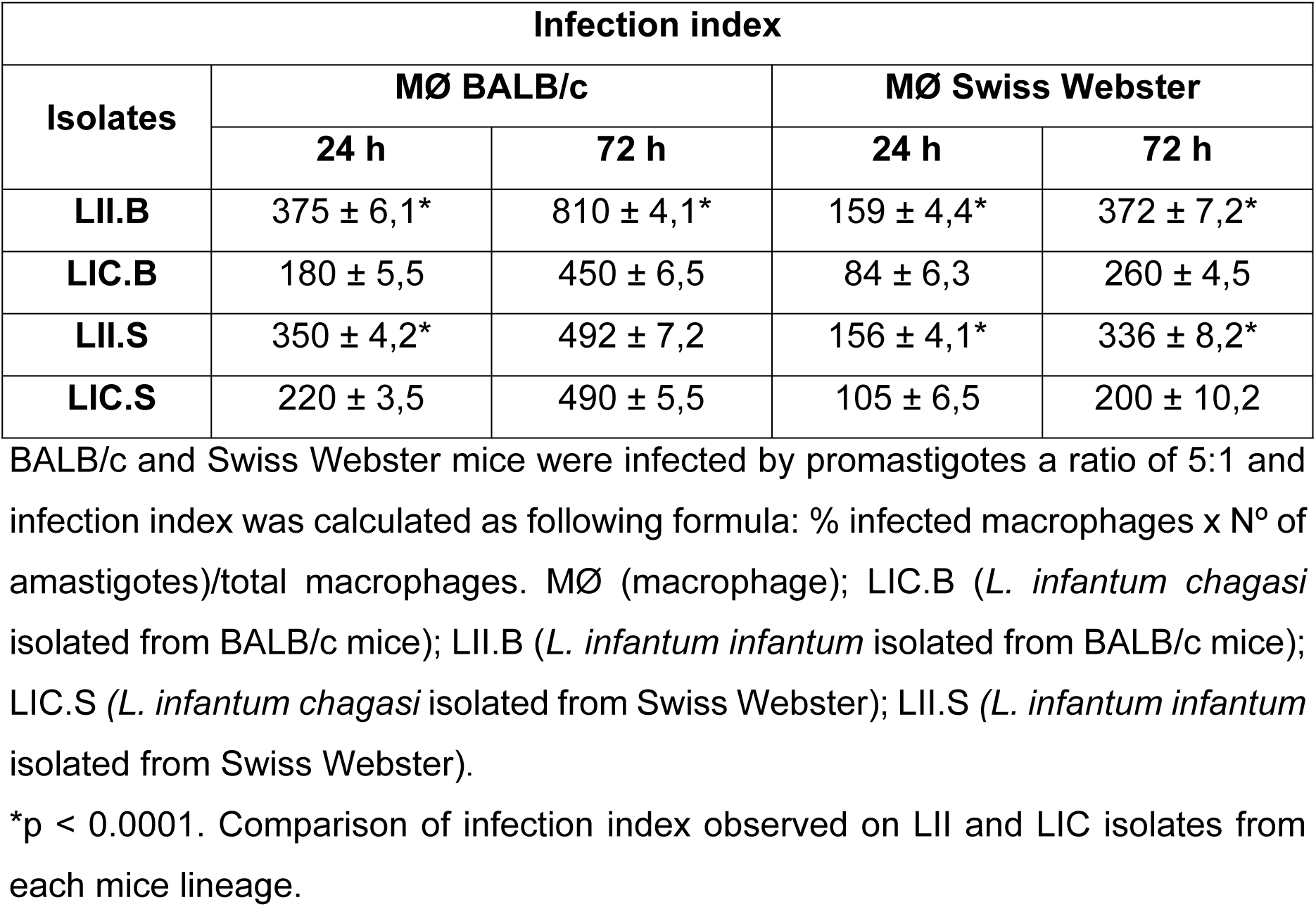
Infectivity of the four *Leishmania* isolates to peritoneal macrophages.

| Isolates | Infection index |  |  |  |
| --- | --- | --- | --- | --- |
|  | MØ BALB/c |  | MØ Swiss Webster |  |
|  | 24 h | 72 h | 24 h | 72 h |
| **LII.B** | 375 ± 6,1* | 810 ± 4,1* | 159 ± 4,4* | 372 ± 7,2* |
| **LIC.B** | 180 ± 5,5 | 450 ± 6,5 | 84 ± 6,3 | 260 ± 4,5 |
| **LII.S** | 350 ± 4,2* | 492 ± 7,2 | 156 ± 4,1* | 336 ± 8,2* |
| **LIC.S** | 220 ± 3,5 | 490 ± 5,5 | 105 ± 6,5 | 200 ± 10,2 |

BALB/c and Swiss Webster mice were infected by promastigotes a ratio of 5:1 and infection index was calculated as following formula: % infected macrophages x N° of amastigotes)/total macrophages. MØ (macrophage); LIC.B (*L. infantum chagasi* isolated from BALB/c mice); LII.B (*L. infantum infantum* isolated from BALB/c mice); LIC.S (*L. infantum chagasi* isolated from Swiss Webster); LII.S (*L. infantum infantum* isolated from Swiss Webster).
\*p < 0.0001. Comparison of infection index observed on LII and LIC isolates from each mice lineage.

### NOS activity

NOS activity was evaluated in promastigotes derived from amastigotes isolated from the four experimental groups, i.e. LII.B, LIC.B, LII.S and LIC.S. Interestingly, the intracellular NO production in the four isolates showed no significant differences, possibly because LII and LIC produce similar basal levels of NO (Fig 4A).

**Figure 4.**
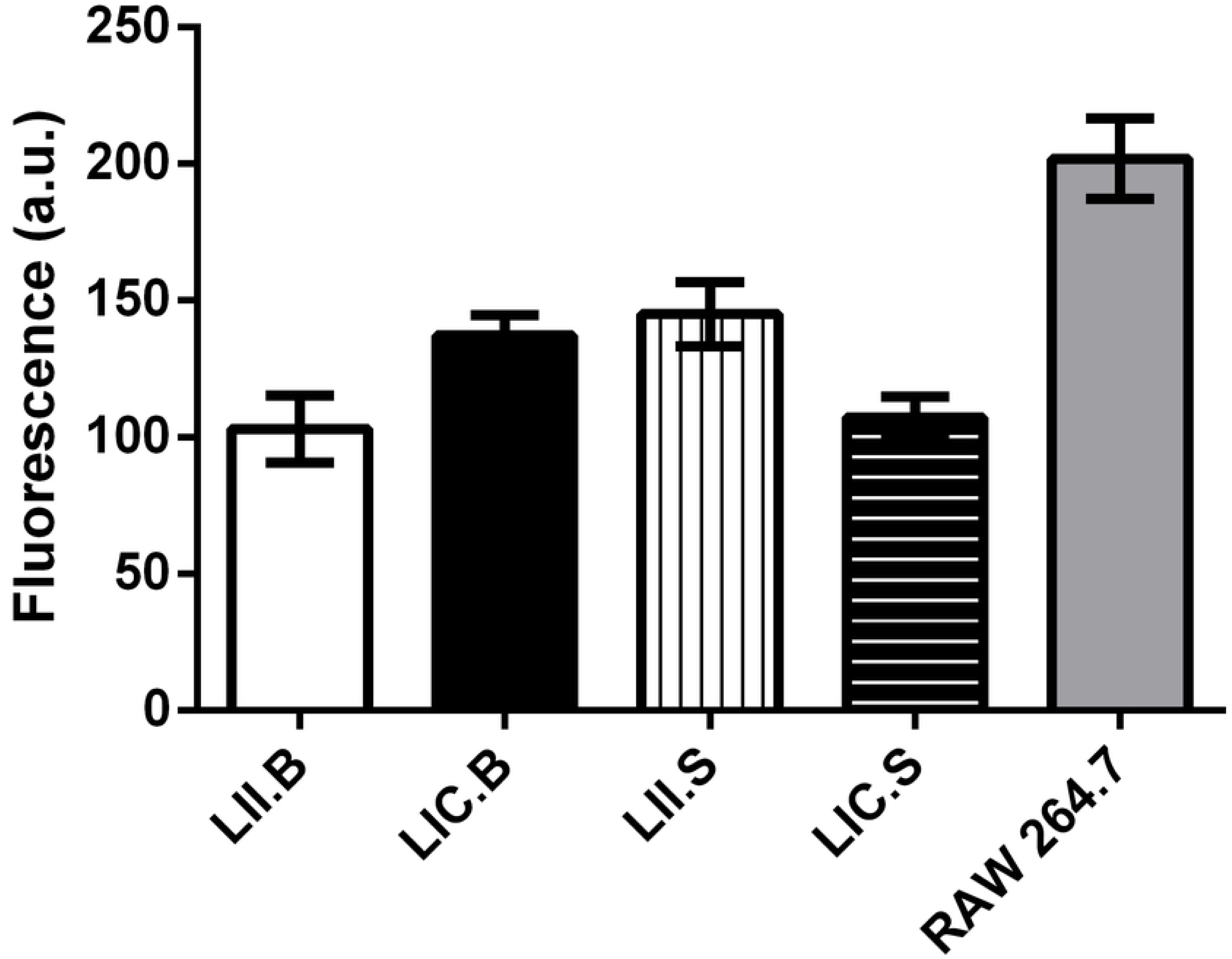
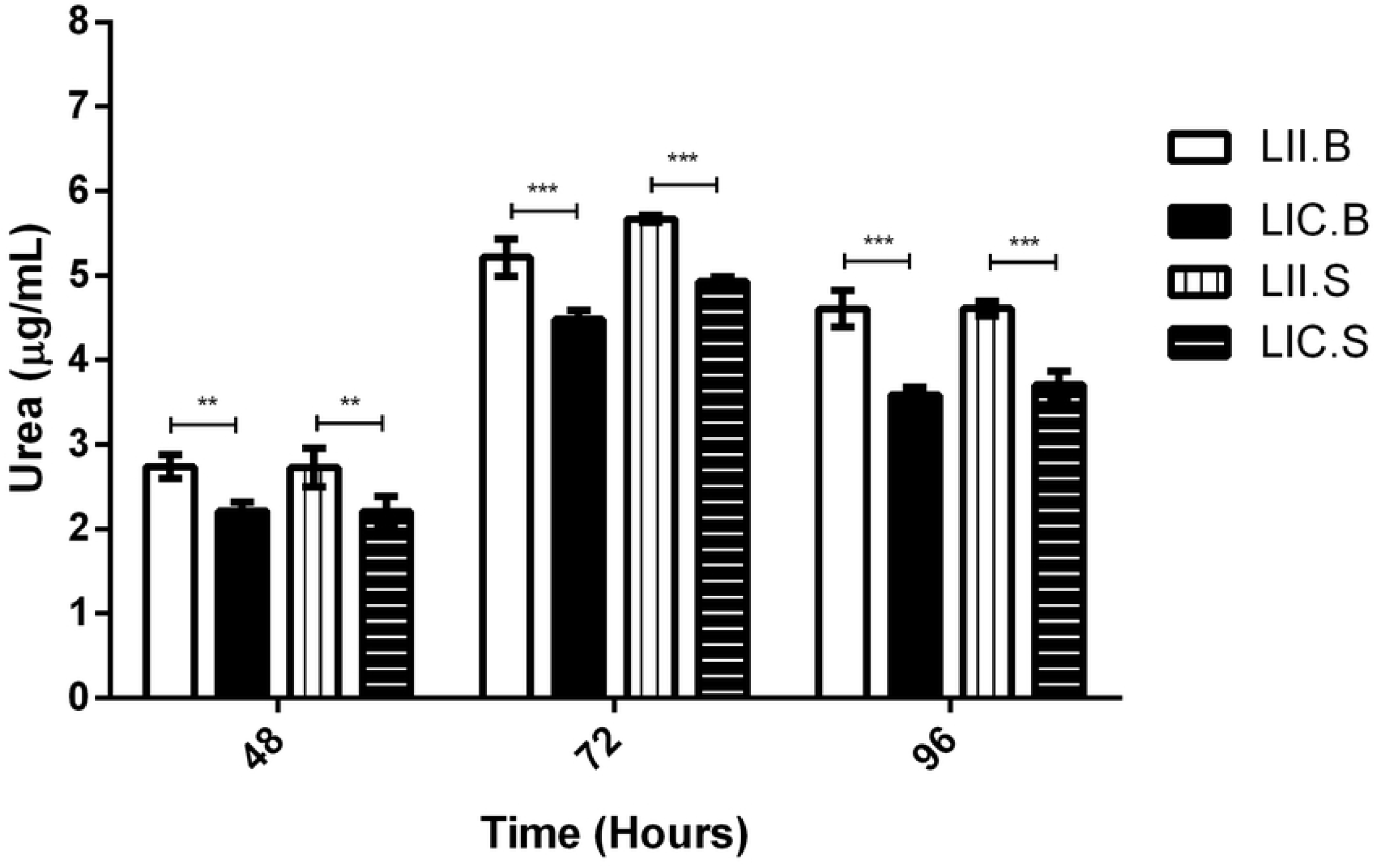
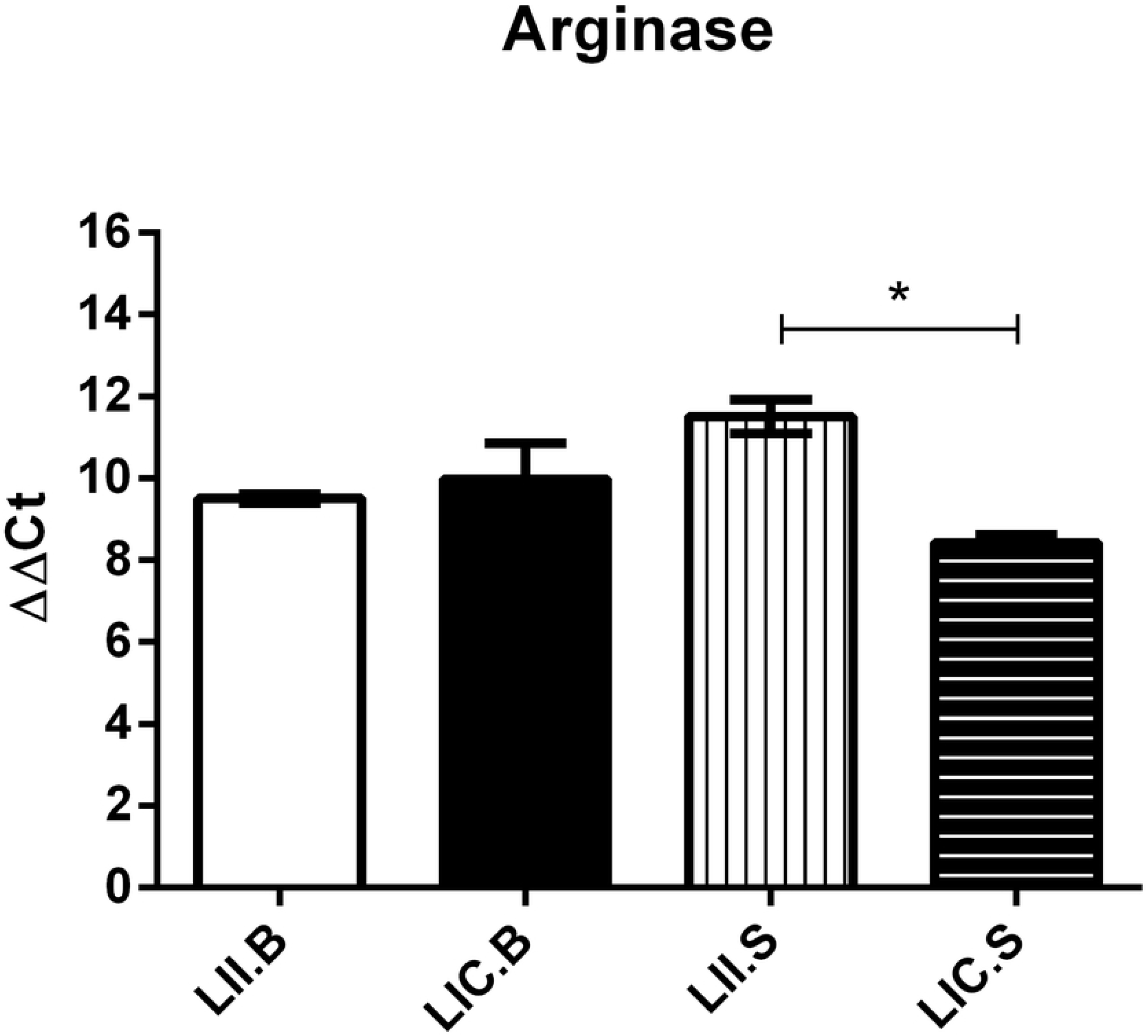
*In vitro* NOS and ARG activities. (A) NOS activity of promastigotes. (B) ARG activity by urea production by promastigotes. (C) Relative expression of arginase gene. The intracellular NO production was detected by DAF-2DA labelling. RAW 264.7 macrophages were used as positive control. The urea production was measured by spectrometry at 540 nm. The relative expression of arginase gene was evaluated by qRT-PCR. alfa-Tubulin gene was use as reference gene. LIC.B (*L. infantum chagasi* isolated from BALB/c mice); LII.B (*L. infantum infantum* isolated from BALB/c mice); LIC.S *(L. infantum chagasi* isolated from Swiss Webster); LII.S *(L. infantum infantum* isolated from Swiss Webster). *p ≤ 0.05; **p ≤ 0.009; ***p < 0.0001

### ARG activity and gene expression

ARG activity in promastigotes of the four isolates was measured after 48 to 96 h in culture, and higher enzyme levels were obtained in 72 h of cultures. However, in all points evaluated, LII isolated from both mice lineages (LII.B and LII.S) showed higher urea levels compared to LIC, demonstrating highest ARG activity (Fig 4B). Higher enzyme levels occurred in LII parasites with higher percentages of metacyclic forms, suggesting that ARG activity may be related to the parasite infectivity. Since both LIC and LII showed differences in the *in vivo* ARG activity, the expression of this enzyme was evaluated in the four *Leishmania* isolates. As observed, LII.S parasites (*L. infantum infantum* isolated from Swiss Webster) showed an increase (p ≤ 0.05) in ARG expression compared to LIC.S (*L. infantum chagasi* isolated from Swiss Webster) (Fig 4C), indicating that such differences may be associated with the enzyme gene expression. To confirm this hypothesis, genetic sequencing was performed. However, analysing the ARG sequences of the four *Leishmania* isolates, no differences were observed between the sequences of LII and LIC. Thus, it is possible to state that the strains are molecularly equal (relative to the ARG gene) and identical to the sequence of *L. infantum* deposited in the TriTrypDB database.

## Discussion and Conclusions

Since VL is a tropical disease that affects millions of people and can lead to death, the study of this pathology, as well as the understanding of the characteristics of its etiological agents, is of fundamental importance. As previously reported, the classification of the parasite responsible for American Visceral Leishmaniasis (AVL) is still controversial. Considering *L. infantum chagasi* (LIC) as a different species, or as a synonymous of *L. infantum infantum* (LII), should take into account the biological, biochemical and pathogenic behaviours ^[27]^. In the literature, there are several studies demonstrating specific genetic and molecular variations among *L. infantum* isolates from different geographic regions ^[28]^, however, few studies address the biological differences, as well as the behaviour of these strains during infection. In this work, the *in vitro* and *in vivo* differences in the infectivity and survival capacity of LIC and LII were reported.

It was observed that these isolates showed different infectivity levels in experimental infection *in vivo* in two different murine models. Genetic factors dependent on the mice lineage interfere with the success of the experimental infection, leading to differences in disease development. Thus, the choice of an animal model is of great relevance, since the genetic background influences both immunological and pharmacological responses to chemotherapeutic agents. Also, during infection by *Leishmania* spp., the tropism of each parasite species in the vertebrate host, must be consider when planning the experimental design.

It is already established that different elements of the host immune response directly influence the course of the infection ^[29,30]^. It is known that T-helper cells-mediated response plays a key role in the control of infection by *Leishmania* parasites. Such immune response is clearly polarized in murine models, with resistant animals showing a higher production of INF-γ, whereas susceptible ones, increased IL-4 production ^[31]^. The Th1 and Th2 polarization is determined by environmental and genetic factors, explaining different susceptibilities to infection in mice with different genetic backgrounds. In the present study, BALB/c mice were more susceptible to *Leishmania* infantum infection than Swiss Webster mice.

It is already known that parasites of the genus *Leishmania* have the ability to modulate the immune response of the infected host to increase their chances of intracellular survival ^[32]^. One of these mechanisms is the uptake of L-arginine, a substrate used by the enzyme iNOS, to suppress NO production in the host and inhibit the release of pro-inflammatory cytokines ^[33]^. This process leads to an increase in ARG activity, which in turn can be modulated by the parasite. *In vitro* studies of L-arginine addition to the culture medium showed that the concentration of this amino acid is crucial in the effect of NO and thus on parasite death ^[7]^.

The data here obtained demonstrates that spleen and liver cells from infected mice produced different levels of NO, with the highest levels observed in BALB/c cells infected with LIC. Interestingly, although infected spleen cells produced NO more than uninfected ones, both LIC and LII were still able to survive and multiply within the host, increasing parasite load at 60 dpi, especially in the case of LII. Marques and collaborators (2015) ^[34]^ have reported that LII parasites were able to survive high amounts of NO added to the culture.

Regarding ARG activity, lower urea production was observed in both spleen and liver of infected animals when compared to uninfected controls. However, spleen cells from group BALB/c infected by LII showed higher urea production than of the other infected groups. This higher ARG production may be correlated to higher parasite loads observed in spleen cells. Taken together, the results of the NOS and ARG activities suggest that *L. infantum* parasites may influence the metabolism of L-arginine from the infected host.

In addition to the host’s ability to respond to infection, the biological differences between the parasite strains and parasite species even been expected, should be considered. As published studies have shown that isolates of *L. infantum* belonging to the same zymodeme, which may present different sensitivities to the host’s immune response, which may be related to parasite virulence factors ^[27]^. There are studies indicating that differences in infectivity between *L. infantum* strains could be related to modulation by the parasites of the expression of L-arginine metabolic enzymes during mouse infection ^[34]^. Studies about differences between parasites from different geographic regions are needed. Therefore, the present study aimed to evaluate the biological and molecular characteristics of different isolates of *L. infantum* and their behaviour when isolated from different hosts (LIC.B, LII.B, LIC.S and LII.S). Both LII isolates (LII.B as LII.S) showed a higher growth rate than LIC.B and LIC.S, even considering that all the four isolates present the same proliferation profile. The infectivity and behaviour of different parasites during macrophage infection from the same hosts from which they were isolated were also evaluated. As in the *in vivo* results, LII was shown to be more infective both by the percentage of metacyclic forms in culture, as well as by the infection index in murine macrophages. In addition, macrophages from mice with different genetic backgrounds also presented different susceptibilities to *in vitro* infection, reinforcing the importance of the search for better murine models for the experimental infection.

*Leishmania* parasites when established within macrophages demonstrate an ability to survive in a “hostile” environment and need to modulate the host immune response. L-Arginine metabolism is closely associated with the success/failure of the establishment of the *Leishmania* spp. infection, since this route can determine death/survival of the parasite ^[15]^.

In this sense, the decrease in NO production in *Leishmania*-infected macrophages has been demonstrated by the inhibition of the iNOS enzyme ^[36]^. Studies with *L. donovani* showed the ability of the parasite to increase ARG activity in spleen cells, favouring the establishment of infection ^[37]^. It is therefore of great interest to establish a relationship between the levels of induction and inhibition of NOS and ARG in intracellular parasites, as well as in the infected host cell itself. However, few studies have reported differences in the activity of L-arginine metabolic enzymes, mainly comparing promastigotes from different strains of *L. infantum*.

NO production by promastigotes of the four isolates was determined by Green’s method, however, due to the low amounts of nitrite present in the culture supernatants, this technique was not sensitive enough. So, the fluorescent indicator DAF-2DA was employed, allowing the detection of basal NO levels and no significant differences were observed among the four isolates.

As demonstrated by Balestieri and collaborators (2002) ^[36]^, inhibition of iNOS activity was observed in *Leishmania*-infected macrophages followed by reduction of NO production, indicating the parasite’s ability to evade host cell defence mechanisms. Although NO is a molecule responsible for several cellular signalling processes and is considered as one of the key pieces in the host’s immune response ^[37]^, ARG is equally important, since it is an essential enzyme in the polyamine pathway. However, few papers in the literature report the importance of parasite ARG inhibition and NO production by infected macrophages ^[38]^, even less the activity and expression of both enzymes in different strains of *Leishmania*.

The existence of NOS in *Leishmania* spp. ^[6]^, as well as in *Trypanosoma cruzi* ^[39]^, has already been reported. However, the identification of the gene responsible for encoding this enzyme in trypanosomatids ^[40]^ is insufficient in the literature and in genomic databases, making it difficult to search for molecular differences in NOS among *Leishmania* strains. However, the ARG gene is already well described, thus allowing several analyses to be performed. It was demonstrated in this work that different isolates of *L. infantum* present different ARG activities during their proliferation. However, both LII and LIC isolates showed higher enzyme activity on the third day of culture, when parasites are reaching the stationary phase. To search possible differences in ARG gene sequencing and in the relative expression of the enzyme, sequencing and expression assays were performed. As a result, it was found that ARG gene is identical in LIC and LII and identical to the sequence deposited in the online database TriTryp (data not shown). On the other hand, the four isolates presented comparable differentiated mRNA expression relative to each other.

The data presented in this study indicate that the strains of *L. infantum* from the Old and New World present different biological behaviours, although they are considered identical by some authors. This demonstrates the importance of conducting further studies on these differences, as this may assist in the search for better alternatives to combat these parasites.

It was also concluded that mice with different genetic backgrounds present different susceptibilities to infection by *Leishmania* strains, both *in vivo* and *in vitro*, reinforcing the importance of the search for better animal models to study the pathophysiology of leishmaniases. In addition, L-Arginine metabolism in these parasites can be seen as a good therapeutic target, considering that NO synthase and arginase enzymes can modulate host response to infection.

## Conflict of interest

The author(s) declare(s) that they have no conflict of interest to disclose

## Acknowledgements

We are grateful to the support given by Conselho Nacional de Desenvolvimento Científico e Tecnológico (CNPq), and Fundação Oswaldo Cruz (Fiocruz).

## Supporting information

**Figure S1. Evaluation of *in vivo* infection of two *L. infantum* strains.** (A) Parasite load in the liver of infected mice. (B) Nitrite levels (NOS activity) of liver cell cultures. (C) Urea levels (ARG activity) of liver cell cultures. The BALB/c and Swiss Webster mice infection was maintained at 30 and 30 dpi. The number of parasites/g of liver was estimated based on total weight of the liver removed and the parasite load in the serial dilution. The nitrite and urea levels were measured by spectrophotometry at 540 nm. Control: non-infected mice; LII (*L. infantum infantum*); LIC (*L. infantum chagasi*). *p ≤ 0.05; **p ≤ 0.009; ***p ≤ 0.0009; ***p < 0.0001

